# Local adaptation mediated niche expansion in correlation with genetic richness

**DOI:** 10.1101/2021.11.22.469607

**Authors:** Masaomi Kurokawa, Issei Nishimura, Bei-Wen Ying

**Affiliations:** School of Life and Environmental Sciences, University of Tsukuba, 1-1-1 Tennoudai, Tsukuba, 305-8572 Ibaraki, Japan

**Keywords:** genetic richness, local adaptation, genome reduction, niche expansion, environmental gradient, growth rate, experimental evolution, fitness landscape

## Abstract

As a central issue in evolution and ecology, the quantitative relationship among the genome, adaptation and the niche was investigated. Local adaptation of five *Escherichia coli* strains carrying either the wild-type genome or reduced genomes was achieved by experimental evolution. A high-throughput fitness assay of the ancestor and evolved populations across an environmental gradient of eight niches resulted in a total of 80 fitness curves generated from 2,220 growth curves. Further analyses showed that the increases in both local adaptiveness and niche broadness were negatively correlated with genetic richness. Local adaptation caused common niche expansion, whereas niche expansion for generality or speciality was decided by genetic richness. The order of the mutations accumulated stepwise was correlated with the magnitude of the fitness increase attributed to mutation accumulation. Pre-adaptation probably participated in coordination among genetic richness, local adaptation and niche expansion.

## Introduction

In nature, microorganisms of various genome sizes inhabit a range of environments, i.e., niches, which is the consequence of local adaptation^1^ and is constrained by evolutionary costs^2^. Genome size, i.e., genetic richness, was believed to be associated with the ecological niche^3^, which was supported by the linkages between genome streaming and niche partitioning^4^, gene loss and niche shift^5^, genome reduction and habitat transition^6^ or metabolic cost^7^, and genome architecture and habitat^8^ or niche-directed evolution^9^. These ecological findings and genomic analyses provided strong evidence of the relationship between the genome and niche established during adaptive evolution in nature. However, the quantitative evaluation of this relationship is largely insufficient and might require experimental demonstration in the laboratory.

First, the contribution of genetic richness to adaptative evolution, i.e., local adaptation, remains unclear, although changes in genome size are commonly observed in nature^10,11,12^ and known as the major driving force for adaptive evolution, e.g., horizontal gene transfer^13,14^. Genome size, i.e., genetic richness, was experimentally reduced^15,16,17^ to achieve the minimal genetic requirement for living organisms^18,19^. The reduced genomes tended to show decreased fitness^20,21^ and increased mutagenesis^22^, which could both be restored by experimental evolution^22^. These studies verified the connection between genetic richness and adaptive evolution, but quantitative evaluation via comparison to the wild-type genome was lacking.

Second, the contribution of the local adaptation achieved by evolution to niche broadness was unknown. To date, experimental studies have generally focused on the target component out of numerous components that comprise the environment, e.g., the carbon source^23^ or antibiotics^24^, as the trigger factor for adaptative evolution. The environment, either the culture medium used in the laboratory or the ecological niche in the wild nature, is comprised of not only the target component but also a number of other nutrients and trace elements. The participation of components other than the target component in adaptative evolution was generally neglected. Machine learning predicted that the priorities of the medium components were differentiated in deciding the bacterial growth^25^, indicating the varied adaptiveness in response to the individual components comprised of the environment. The fitness landscape across the environmental gradients of all related components (niches) was crucial for us to address the question of how local adaptation contributes to niche broadness.

The present study addressed the questions of how genetic richness (genome reduction) contributed to local adaptation and whether and how local adaptation caused changes in niche broadness (Fig. 1A). Genetic richness was represented by genome reduction; local adaptation was achieved by experimental evolution; and niche broadness was evaluated by fitness curve fitting across the environmental gradients of the components (niches) presented in the experimental evolution (Fig. 1B).

**Figure 1.**
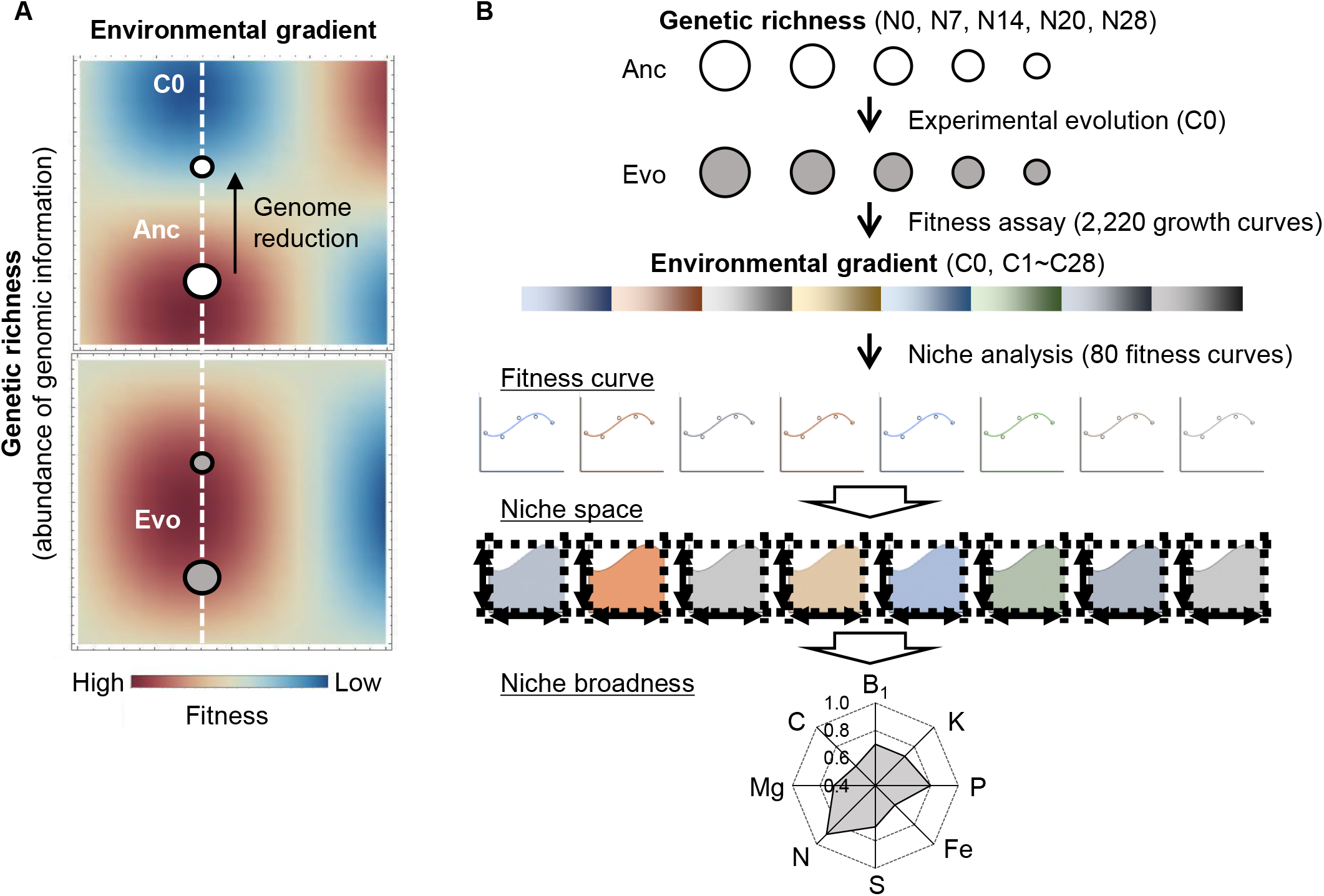
Conceptual illustration of the study. **A.** Fitness landscapes of the reduced genomes in the environmental gradient. Environment. The evolution of the genomes (open circles), from Anc to Evo, leads to not only local adaptation to the environment for evolution (C0, white broken line) but also changes in fitness landscapes across the environmental gradient. The colour gradation from red to blue indicates the fitness from high to low. **B.** Overview of the experiments and analyses performed in the present study. Five genomes (strains) are shown as circles. Black arrows indicate the experimental and/or analytical studies. The keywords newly defined in the present study are underlined. The colour variation and gradation represent the difference in the chemical components (niches) and the concentration gradient, respectively.

## Results

### Experimental evolution-mediated local adaptation was correlated with genetic richness

Local adaptation, which was achieved by experimental evolution with *E. coli* strains of varied genome sizes, showed that the fitness increase was correlated with genetic richness. Five laboratory *E. coli* strains carrying either the wild-type (N0) or the reduced (N7, 14, 20, 28) genomes (Table S1) were subjected to experimental evolution in chemically defined medium (designated C0). A gradual increase in the growth rate was commonly observed in the reduced genomes during evolution for approximately 1,000 generations (Fig. 2A). The growth rates of the evolved populations (Evos) were all higher than those of the ancestors (Ancs), indicating that local adaptation was achieved regardless of genetic richness (Fig. 2B). This finding was somehow consistent with our previous finding that genome reduction was correlated with a decrease in the growth rate^20^.

**Figure 2.**
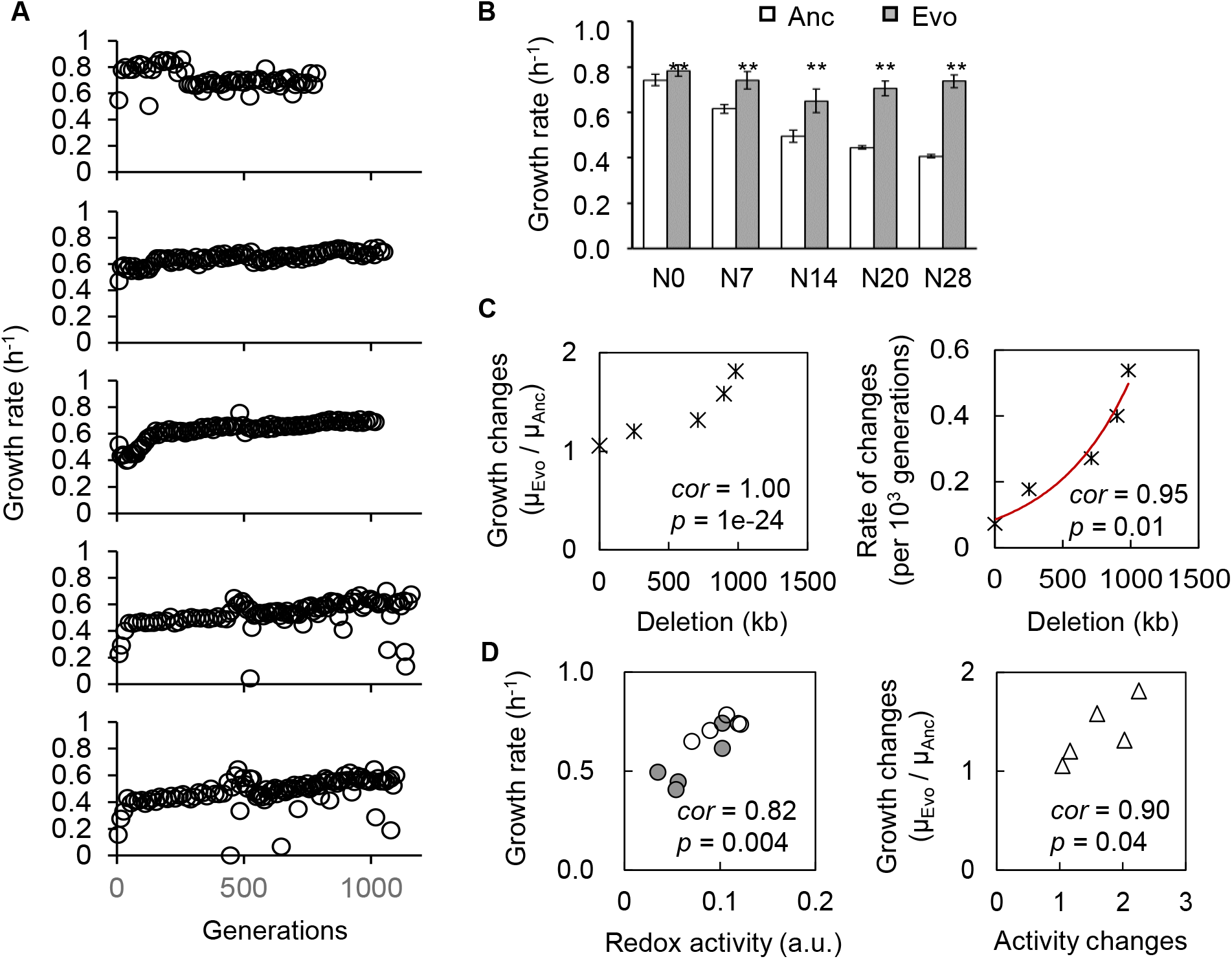
Correlation between local adaptation and genome reduction. **A.** Temporal changes in growth rates during the experimental evolution. The five genomes are indicated. **B.** Growth rates in the medium for evolution. Open and closed bars represent the growth rates of Ancs and Evos in the medium C0, respectively. Standard errors of biological replications (n > 6) are indicated. Asterisks indicate the statistical significance of the two-tailed Student’s t-test (*p* < 0.01). **C.** Correlation between growth changes and genome reduction. The lengths of the genomic deletions are plotted against the ratio of the growth rates of Anc and Evo (left) and the ratio per generation (right). The Spearman rank correlation coefficients and statistical significance are indicated. The red line indicates the logarithmic regression. **D.** Correlation between the growth rate and cellular redox activity. The left and right panels show the relationships between the growth rate and the redox activity (NADH) and the changes in both, respectively. Open and closed circles represent Ancs and Evos, respectively. The Spearman rank correlation coefficients and statistical significance are indicated.

Both the changes in growth rate and the rates of the changes in evolution were significantly correlated with genome reduction (Fig. 2C), indicating coordination between local adaptation and genetic richness. The correlation of genome reduction with the evolutionary rate of fitness increase was supported by the previous finding of the correlation between genome reduction and the spontaneous mutation rate^22^, a global parameter representing evolvability. Additionally, the cellular redox activity, representing the metabolic activity inside the cell, was found to be correlated with the growth rate, and the changes in growth rate and the changes in redox activity were correlated (Fig. 2D). The results further verified that local adaptation was achieved metabolically.

### Local adaptation caused niche expansion in correlation with genetic richness

Whether the local adaptation caused the fitness changes across the environmental gradient was analysed. The growth fitness of Ancs and Evos was precisely evaluated in 29 medium combinations (C0 and C1~28), which comprised eight chemical components used for the evolutionary condition C0 (Fig. 3A, Table S2). A total of 2,220 growth rates calculated from the corresponding growth curves were acquired (Table S5). A global increase in growth rate under most medium combinations was observed in the reduced genomes (Fig. 3B, Fig. S2). Fitness improvement was achieved not only under evolutionary condition C0 but also across the concentration gradient. It seemed that larger deletions from the genome led to larger changes in growth fitness. In comparison, the growth rates of the wild-type genome (N0) were slightly changed. The results revealed that whether local adaptation triggered global adaptation across the environmental gradient was dependent on genetic richness.

**Figure 3.**
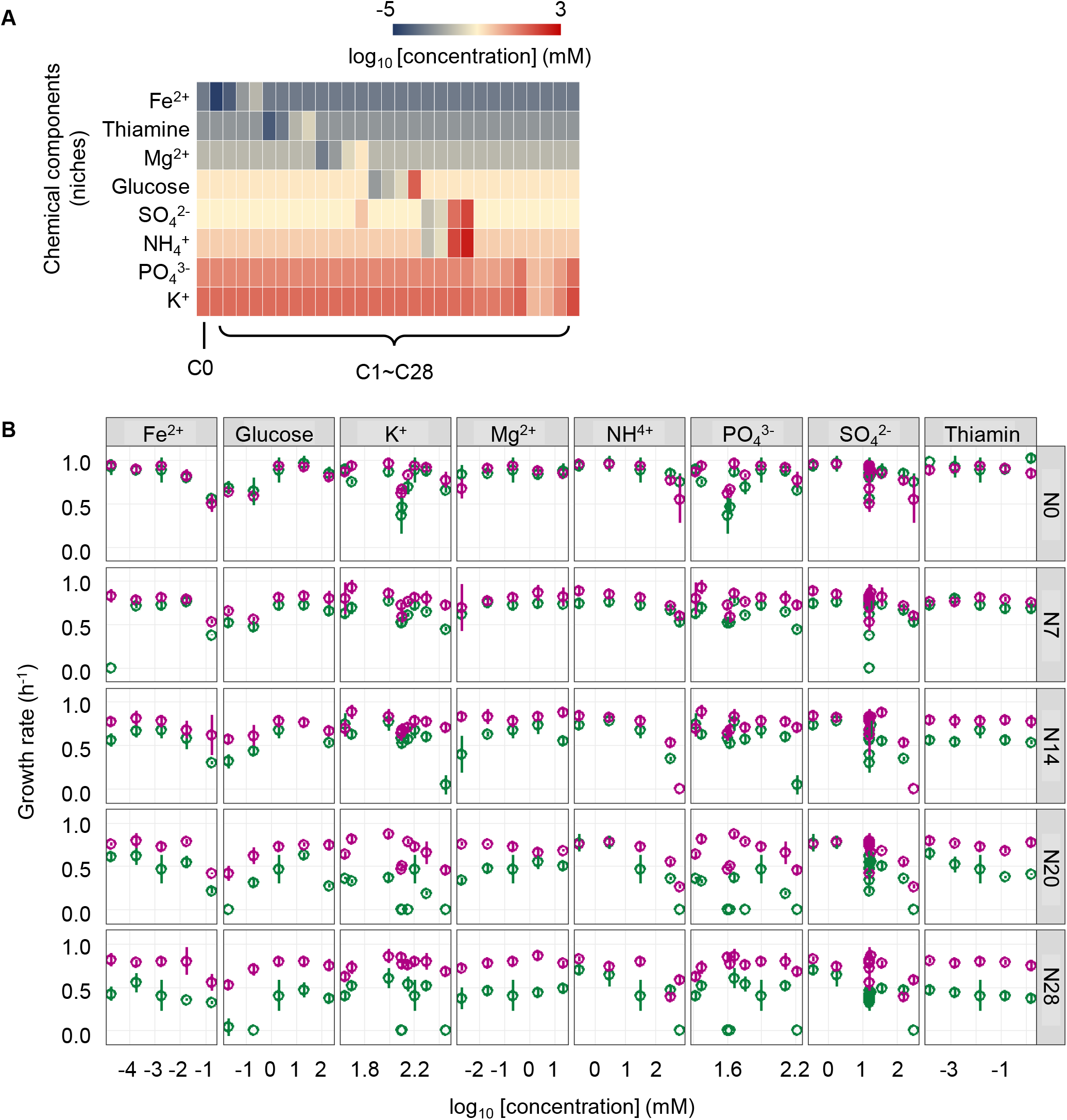
Growth fitness across the environmental gradient. **A.** Heatmap of the concentration gradient. The chemical components of the media are indicated. The concentrations are shown on the logarithmic scale. C0 is the environment for evolution. Both C0 and C1~C28 were used for the fitness assay. **B.** Growth fitness across the concentration gradient of individual chemical components. The mean growth rates in the 29 medium combinations are shown. The concentrations of the chemical components are shown on a logarithmic scale. The wild-type and reduced genomes are indicated as N0 and N7~N28, respectively. Purple and green represent Evo and Anc, respectively. Standard errors of biological replications (n > 6) are indicated.

According to the growth rates (Fig. 3B) in eight chemical components (niches), the niche space (***S***) was newly defined by cubic polynomial regression to the normalized fitness curve, in which the maxima of both the concentration gradient and the growth rate were normalized to one unit (Fig. 4A). The normalization of individual fitness curves determined a fixed niche breath available for the comparison among the varied niches and genomes. The niche space (***S***) was determined as the shadowed space under the regression curve. Consequently, a total of 80 niche spaces (Fig. S3), as well as the changes in niche space attributed to local adaptation (Fig. S4), were calculated. It seemed that both the niche space and its change were more closely associated with genetic richness (genomes) than with the niche types (chemical components). To achieve an overall evaluation of niches, the niche broadness (total ***S***) was defined as the sum of the eight niche spaces (Fig. 4B).

**Figure 4.**
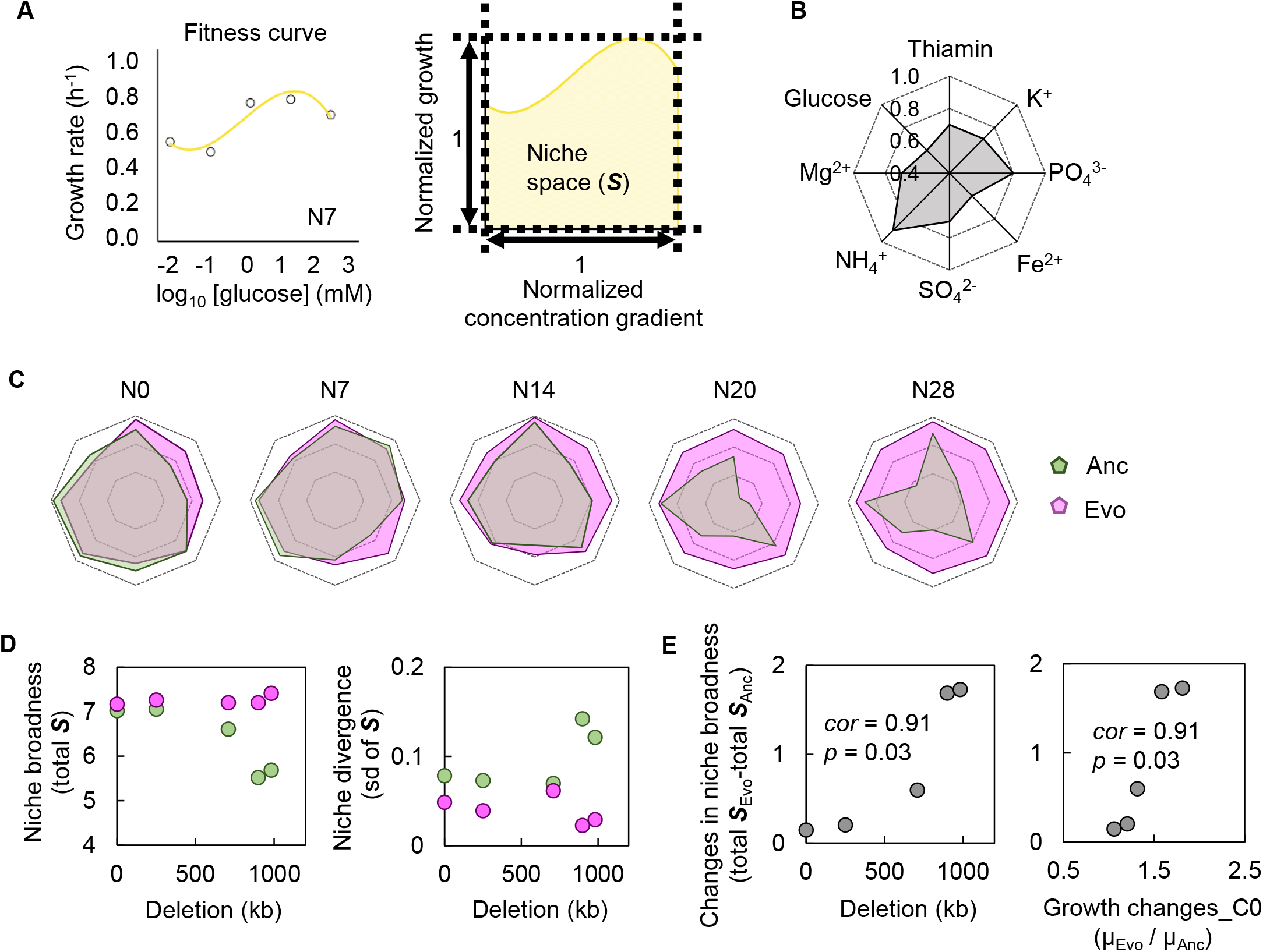
Niche broadness. **A.** Definition of the niche space. The left and right panels indicate the fitness curve of N7 across the concentration gradient of glucose and the B-spline regression of the normalized fitness curve, in which both the concentration gradient and the growth rates are rescaled within one unit, respectively. The area in shadow was determined as the niche space (***S***) of N7 in the niche of glucose. **B.** Radar chart of the eight niche spaces. The niche names, i.e., the eight chemical components, and the scale of the niche space are illustrated in the monotone radar chart. **C.** Evolutionary changes in niche broadness. The five genomes are shown separately. Purple and green represent Evo and Anc, respectively. **D.** Niche broadness and divergence. The sum of the eight niche spaces is defined as the total niche broadness (left). The standard deviation of the eight niche spaces is defined as the niche divergence (right). **E.** Correlation of niche broadness to local adaptation. The changes in niche broadness between Ancs and Evos are plotted against the lengths of the genomic deletion (left) and the changes in growth rates of the local adaptation (right). The Spearman rank correlation coefficients and statistical significance are indicated.

The niche broadness was narrowed in response to genome reduction (Fig. 4C, green); however, it was significantly widened due to experimental evolution (Fig. 4C, pink). The local adaptation expanded the niche broadness of all Evos to a roughly equivalent level (Fig. 4D, left), indicating homeostasis in niche expansion. The variation in spaces of the eight niches commonly declined in the Evos (Fig. 4D, right), indicating that local adaptation reduced niche divergence for balanced niche expansion. Intriguingly, the changes in niche broadness were positively correlated with both genome reduction (Fig. 4E, left) and changes in the growth rate for local adaptation (Fig. 4E, right). These results suggested that local adaptation mediated global coordination of niche expansion with genetic richness for homeostatic and balanced niche broadness.

### Niche expansion for speciality or generality was dependent on the genetic richness

A gradual shift from the evolutionary trade-off in niche space to global niche expansion occurred in response to genome reduction (Fig. 4C). The niche broadness of Anc and Evo entirely overlapped in the reduced genomes with larger deletions (N14, N20, N28) but partially overlapped in the wild-type genome (N0) and the reduced genome with a relatively small deletion (N7). For instance, trade-offs occurred in N0; that is, the improvement in the niches of thiamin, K^+^, PO_4_^+^ and Fe^2+^ and the deficiency in the niches of glucose, Mg^2+^, NH_4_^+^ and SO_4_^2+^ implied niche expansion for speciality. In contrast, omnidirectional expansion of niche broadness occurred in the reduced genomes of N14, N20 and N28. These results revealed that whether local adaptation triggered global adaptation across the environmental gradient or the niche-dependent trade-off was dependent on the genetic richness.

A niche-specific correlation of the changes in niche space to local adaptation and genome reduction was observed (Fig. 5). Significant correlations of niche expansion (i.e., changes in niche space) with local adaptation (i.e., changes in growth rate in C0) (Fig. 5A) and genetic richness (i.e., genome reduction) (Fig. 5B) were commonly found in the niches of glucose, SO_4_^2+^ and Mg^2+^. The co-correlation of the local adaptation and the genetic richness to niche expansion was consistent with the correlation between the growth change and genome reduction (Fig. 2C). Note that the trade-off in niche expansion of N0 reduced the niche space of glucose, SO_4_^2+^ and Mg^2+^ (Fig. 4C), in which niche expansion was significantly correlated with genome reduction (Fig. 5B). This implied that the genetic richness was highly sensitive to these three niches with regard to carbon, sulphate and magnesium.

**Figure 5.**
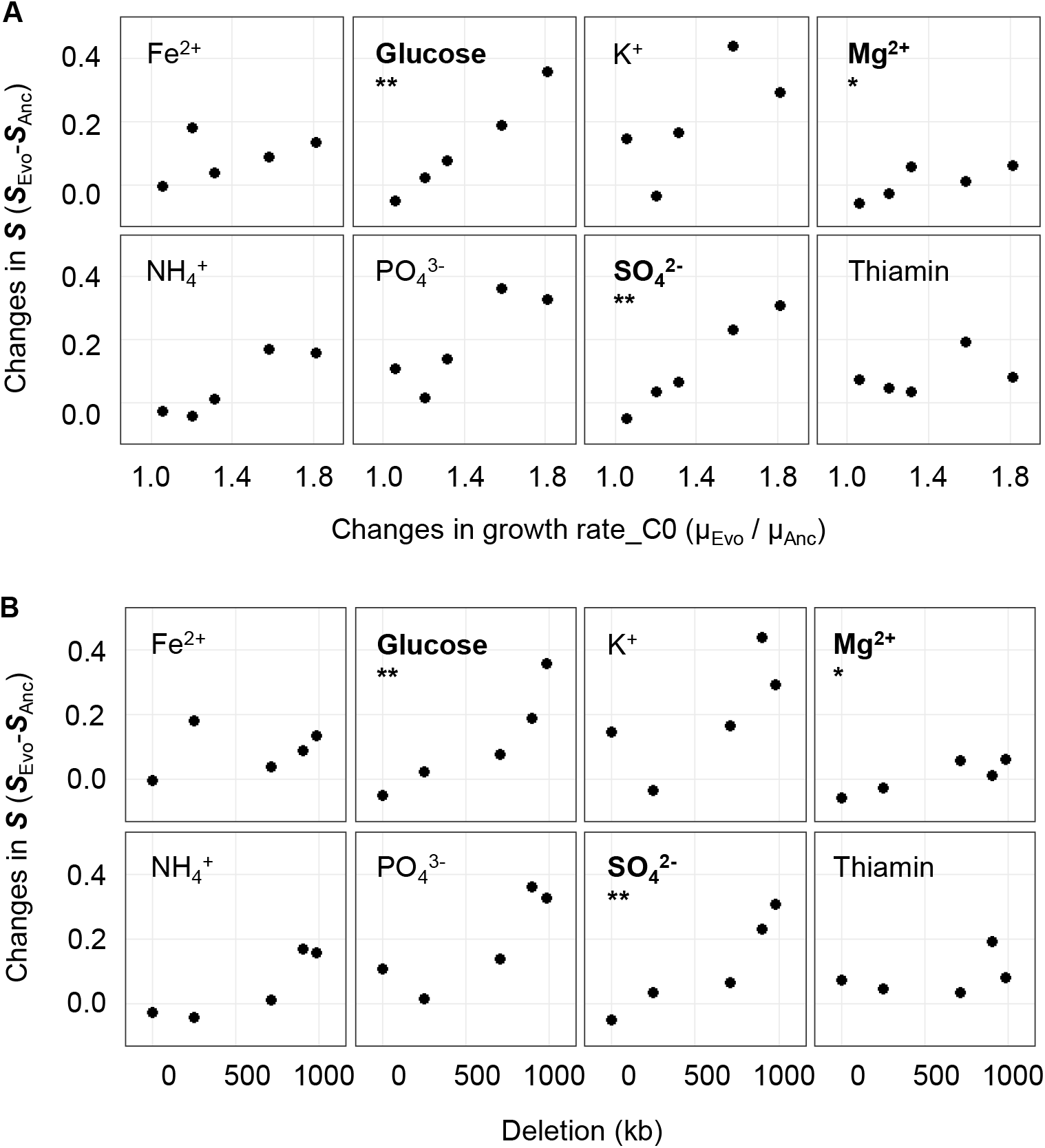
Niche-differentiated changes in the niche space caused by local adaptation. **A.** Relationships between the changes in niche space and the changes in growth rate. The changes in niche space of the five genomes are plotted against the changes in growth rate in the medium C0 with respect to the eight chemical components (niches). **B.** Relationships between the changes in niche space and the lengths of genomic deletion. The chemical components (niches) and the statistical significance of the Spearman rank correlation are indicated. Boldfaces associated with asterisks represent statistical significance (*,*p* < 0.05; **, *p* < 0.01).

### Stepwise mutation accumulation was associated with an additive increase in fitness regardless of genetic richness

To discover the genetic mechanism participating in local adaptation-associated niche expansion, genome mutation analysis was performed. An approximately equivalent number of gene mutations were detected in the Evos, regardless of the genome (Fig. 6A, Table S3). Temporal changes in the allele frequency of mutants showed that the mutations accumulated serially and were fixed in a stepwise manner. In addition, only a few gene mutations were able to compensate for the large genomic deficiency (Fig. 6A), which was somehow consistent with the finding that the mutations caused the metabolic rewiring of a reduced genome due to experimental evolution ^26^. Note that the changes in transposons were ignored, and the mutations fixed in Evos were identified in the reduced genomes but not in the wild-type genome. The mutated genes were related to transporters and regulators (Table S3), which indicated that resource diffusion for utilization and global gene regulation contributed to local adaptation-associated niche expansion.

**Figure 6.**
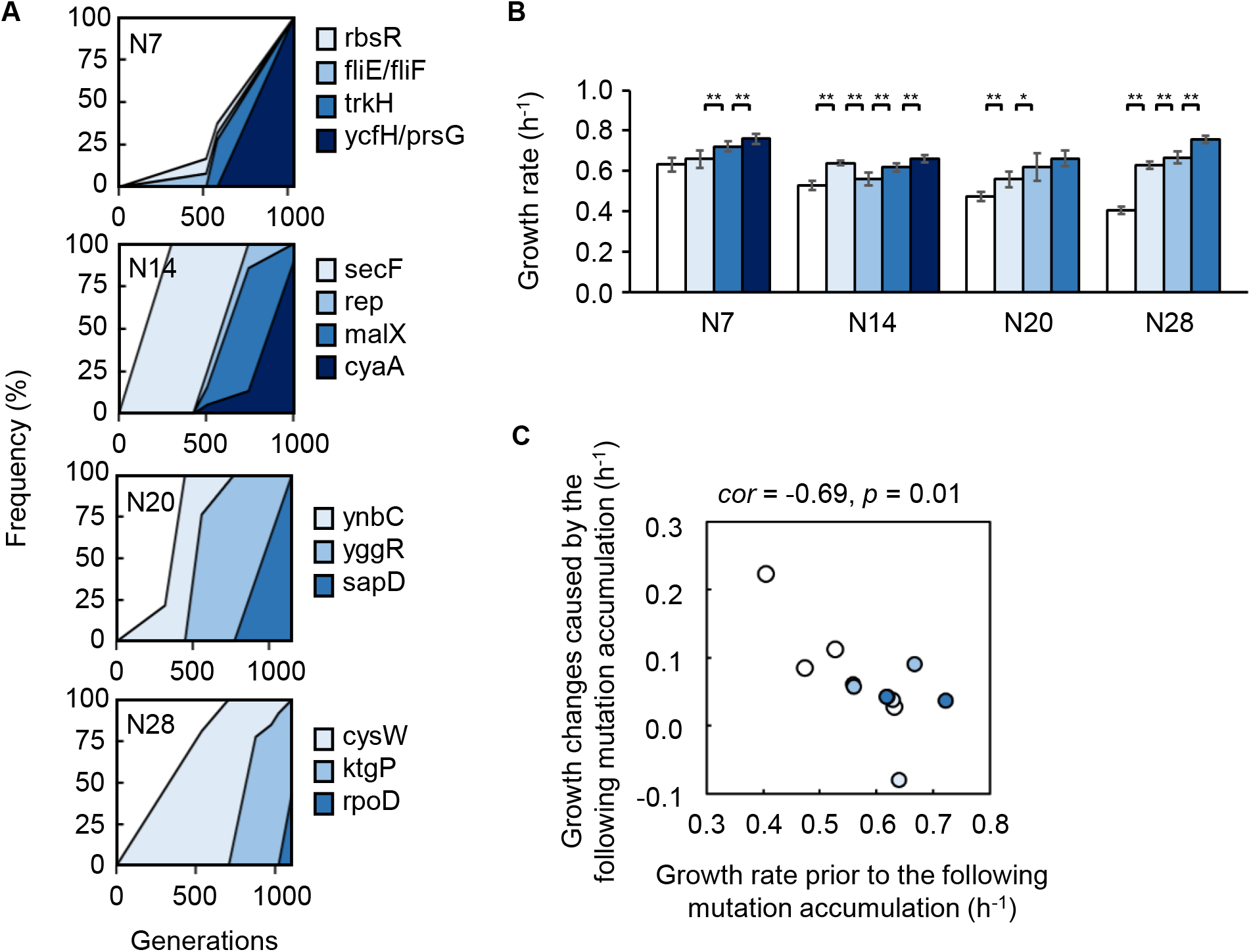
Fitness increase attributed to the mutations. **A.** Stepwise accumulation of genome mutations. Mutation during the evolution. The temporal changes in the mutations fixed during the evolution are shown. The four reduced genomes and the gene names of mutants are indicated. **B.** Additive increase in the growth rates of the mutants in the medium for evolution. Gradation from white to dark blue indicates the mutants with respect to those shown in **A**. Standard errors of biological replications (n > 6) are indicated. Asterisks indicate the statistical significance of the two-tailed Student’s t-test (*, *p* < 0.05; **, *p* < 0.01). **C.** Correlation between mutation accumulation and changes in growth rates. A total of 12 mutants of a single accumulated mutation are shown. The Spearman rank correlation coefficients and statistical significance are indicated.

How the stepwise accumulation of the mutations contributed to local adaptation was further investigated. The colonies/mutants carrying the mutations in the order of evolutionary accumulation were acquired and subjected to a growth fitness assay. Note that the mutants with the second mutation (*rbsR* and *fliE/fliF*) in N7 failed to be acquired, indicating the co-fixation of these two mutations during evolution. A gradual increase in the growth rate of the colonies in the order of mutation accumulation was commonly observed, except for a transient decrease caused by the second mutation that occurred in N14 (Fig. 6B). The mutation accumulation-associated increase in growth rates was commonly observed in all the reduced genomes. This demonstrated that the mutations were beneficial and contributed to local adaptation in an additive manner, which was independent of genetic richness. In addition, an intriguing power law for the contribution of the gene mutations to the local adaptation was observed; that is, lower growth rates prior to mutation fixation led to larger changes in growth rate after the mutation was fixed (Fig. 6C). The first mutations were more likely to improve the growth fitness than the mutations that were fixed later, although the statistical significance was weak because there were too few mutations (Fig. S5). The negative correlation and the order of the fitness contribution of the mutations indicated the pre-adaptation proposed for the evolution of diversity^27^ and the predictivity of the mutation-mediated fitness landscape^28^.

### Hypothesis of pre-adaptation in local adaptation-associated niche expansion

The mechanism linking the fitness landscape, which was commonly applied for explaining evolutionary adaptation^29^, to pre-adaptation for niche expansion was proposed as follows (Fig. 7A). **i**) Growth fitness was correlated with genome reduction^20^. Larger deletions led to a greater distance from the fitness peak. Not only the gene function but also the genetic richness might have played a role in the fitness landscape. Increasing the genome size might have been easier than evolving the well-regulated gene function. **ii**) Evolution improved the local adaptation to C0 in correlation with genetic richness. The greater the distance from the fitness peak was, the larger change it was to reach the equivalent adaptiveness (fitness). This was supported by the increase in growth rate caused by experimental evolution and the correlation between genetic richness and changes in growth rate (Fig. 2). The mutations were fixed in a stepwise manner during evolution, leading to an additive fitness increase (Fig. 6). **iii**) In alternative environments, genetic richness determines whether there is global adaptation for niche generalization or a trade-off for niche specialization, which is supported by the correlation between niche broadness and genome reduction (Fig. 4). The location in the initial fitness landscape (e.g., C0) likely determined the fitness in the alternative environmental gradient (e.g., C_N_). That is, there was a higher probability of trade-off closer to the local adaptive peak, whereas farther from the local adaptive peak, there was more opportunity for global adaptation, which was consistent with the pleiotropic costs for carbon utilization found in the experimental population^30^. The locational bias must have contributed to the pre-adaptation in evolution.

**Figure 7.**
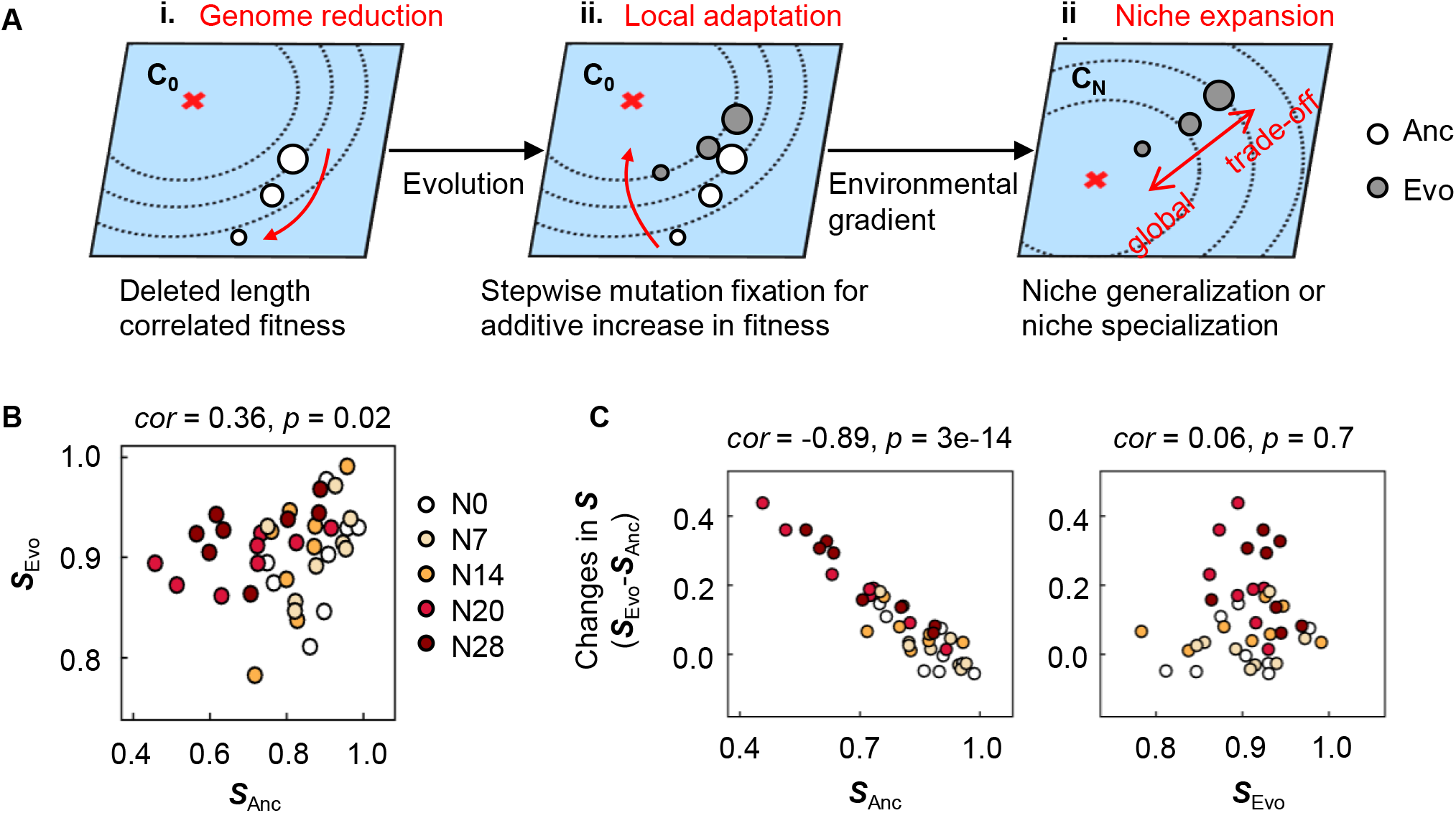
Mechanism for pre-adaptation. **A.** Illustration of the proposed mechanism. The fitness landscapes in environment C_0_ for the evolution and the alternative environments C_N_ are shown as the planes in blue. The red cross and the contour lines indicate the fitness peak (maximum) and the fitness gradient, respectively. The open and closed circles indicate the ancestor and evolved genomes, respectively. The size of the circles indicates the genome size. **B.** Correlated niche spaces of Ancs and Evos. The colour variation from white to dark red represents the five different genomes of N0~N28. The niche spaces of Ancs and Evos are named ***S_Anc_*** and ***S_Evo_***, respectively. **C.** Relationships of ***S_Anc_*** and ***S_Evo_*** to the changes in niche spaces. The changes in the niche space caused by the evolution are plotted against ***S_Anc_*** and ***S_Evo_*** in the left and right panels, respectively. The Spearman rank correlation coefficients and statistical significance are indicated.

The pre-adaptation in the participation of the local adaptation-associated niche expansion was supported by the weak but statistically significant correlation of the niche spaces between Ancs and Evos (Fig. 7B). In particular, the changes in niche space were highly significantly correlated with the niche spaces of Ancs but not with those of Evos (Fig. 7C), which was confirmed even if evolutionary generation was taken into account (Fig. S6). The magnitude of the changes was dependent on the Ancs, that is, the smaller niche space evolved for the larger changes in niche space, which was consistent with the genetic richness-correlated niche expansion (Fig. 4). The correlations identified in the niche space and its changes, as well as the fitness contribution of the mutations, indicated pre-adaptation in evolution.

## Discussion

Larger genomic deletion resulted in not only lower fitness^20^ but also faster evolution (Fig. 2). Rapid adaptation occurred for the reduced genomes, which carried fewer nonessential genes, in comparison to the wild-type genome, which contained a full set of genes. The rapid adaptation of the reduced genome was achieved by a few mutations (Fig. 6), indicating that the large genomic deletions were compensated by a single mutation or a few mutations. Abundant genetic information could be substituted with certain gene functions for equivalent adaptiveness, providing intriguing insight into genetic essentiality. The fitness landscape was employed and preliminarily explained the finding, according to the assumption that both the large deletion and the single mutation were located on the identical fitness landscape (Fig. 7A). Fitness landscape analysis^31,32^ is generally applied to explain mutation occurrence^33,34^ and the resultant mutant, with the changed distance to the fitness peak^35,36^ as evolutionary constraint^33,37^. As genome evolution occurred not only via mutations associated with gene function but also due to large fluctuations in genome size, e.g., horizontal gene transfer^38^ and streamlining^11,39^, introducing the fitness landscape to genome reduction, i.e., changes in genome size as well as changes in gene size, was reasonable. The fitness landscape was consistent with the finding that both the single mutations and the large deletions contributed to the fitness in an additive manner.

Owing to the direct comparison of the wild-type and reduced genomes in adaptive evolution, a genetic richness-dependent evolutionary strategy was first observed. The shift from the adaptive trade-off to global adaption was dependent on genetic richness (Fig. 4). As local adaptation often results in maladaptation to alternative environments^23,40,41^, ecological niche speciation is often explained by adaptive trade-offs^40,42,43,44^. Environmental homogeneity is considered one of the deterministic factors for trade-offs^45^, and environmental fluctuations during evolution are thought to be crucial for global adaptation^46^. Since serial transfer was performed to maintain continuous growth in the early exponential phase (Fig. S1), the evolution was supposed to occur in a steady environment with sufficient resources. Nevertheless, global adaptation instead of the trade-off occurred in the reduced genomes. If it was the deleted genes (gene functions) that had been specifically responsible for the adaptive trade-off, the deleted genomic length would never be correlated with the fitness increase. Thus, it was the genetic richness, i.e., the amount of genomic information, but not the particular gene function, that determined the strategies for trade-off or global adaptation to the environment, e.g., the niche or habitat.

Local adaptation was first considered to be achieved in response to all components of the environment. Evaluation of the fitness landscapes across all components of a wide concentration gradient allowed us to find the common niche expansion related to the adaptive evolution (Fig. 1). The niche expansion was attributed to the fluctuation in the concentration of the components accompanied by bacterial growth, as mentioned previously^47^. The niche divergence (Figs. 4, 5) somehow represented the genetic sensitivity to the components, i.e., the chemical types in the niche. The correlation between the niche space and the genetic richness was significant in the niches of glucose, NH_4_^+^ and SO_4_^2+^, whereas the local adaptation cancelled this correlation for NH_4_^+^ and SO_4_^2+^ and changed the correlation from negative to positive for glucose (Fig. S7). The mutations that occurred for local adaptation largely compensated for the genomic deficiency in using these resources, and this was reasonable because carbon, nitrogen and sulphur are the essential major elements for living organisms on Earth^48^. The genetic richness-correlated changes in niche spaces were decided prior to local adaptation in the niches of glucose and SO_4_^2+^ (Fig. S7), indicating that pre-adaptation more likely occurred in the niches of carbon and sulphate.

The local adaptation caused the large omnidirectional expansion of the niche broadness for the largely deficient genomes in comparison to the small directional expansion of the niche broadness for the complete and few deficient genomes (Fig. 4). Genetic richness was probably associated with omnidirectional fitness across the entire niches of the habitat, which was well supported by the finding that the niche broadness of the ancestor determined the evolved niche broadness (Fig. 7B, C). The habitat, composed of multiple niches, might decide the maximum of the overall niche broadness accessible for evolution. In other words, the overall niche broadness of a defined environment seemed to be homeostatic for a defined species, as the niche expanded until comparable broadness was reached (Fig. 4). Note that the homeostasis of niche space was not biased by normalization. As normalization to one unit was performed individually, the maxima of the overall niche spaces could be differentiated in the respective genomes. Niche expansion might reflect the evolutionary direction for generalists or specialists^2,49^. Deficient and sufficient genetic richness evolved for generality and speciality, respectively, indicating a fundamental principle for genome evolution adaptive to ecological niches.

In summary, the present study provided experimental evidence showing that local adaption mediated niche expansion in correlation with the genome through the combination of genomic and environmental gradients. Despite adaptive evolution in steady environments, deficient genomes evolved in a jack-of-all-trades-and-master-of-all manner, which was theoretically proposed as one of three mechanisms for specialism that is widespread in nature^50^, in comparison to the wild-type genome, which adopted a trade-off mechanism, which was generally explained by constraints in phenotypic space^51,52^. In nature, the trade-off strategy might be more frequent and reasonable for costless adaptation and niche expansion during eco-evolution^2,53,54^. As both the genome and the environment participate in ecological evolution, the coordination among genetic richness, adaptiveness and niche broadness revealed a quantitative linkage of adaptive evolution to ecology.

## Materials and methods

### E. coli strains

A total of five *E. coli* strains with either the wild-type or the reduced genome were used, which were selected from the KHK collection^17^, an *E. coli* collection of reduced genomes (from National BioResource Project, National Institute of Genetics, Shizuoka, Japan). The wild-type and four reduced genomes were derived from *E. coli* W3110 and were assigned as N0 and N7, 14, 20, 28, respectively (Table S1), according to previous studies ^20, 22^.

### Media combinations

The minimal medium M63, equivalent to C0, was used for the experimental evolution for local adaptation. Its chemical composition was described in detail previously^20,55^. The concentration gradient of the components of the M63 medium was prepared just before the fitness assay by mixing the stock solutions of individual chemical compounds, which resulted in 28 alternative medium combinations (C1~28). The stock solutions, that is, 1 M glucose, 0.615 M K_2_HPO_4_, 0.382 M KH_2_PO_4_, 0.203 M MgSO_4_, 0.0152 M thiamin/HCl, 0.0018 M FeSO_4_, and 0.766 M (NH_4_)2SO_4_, were sterilized using a sterile syringe filter with a 0.22-μm pore size hydrophilic PVDF membrane (Merck). The concentrations of most chemical compounds were altered one-by-one on a logarithmic scale to achieve a wide range of environmental gradients, as described previously^25^, which led to a total of 28 combinations (Fig. 3A, Table S2). Both the medium used in the evolution (C0) and the alternative medium combinations (C1~28) were used for the fitness assay. The resultant concentrations of individual components in the ionic form are summarized in Table S2.

### Experimental evolution

The experimental evolution of the five *E. coli* strains was performed within the early exponential phase by serial transfer (Fig. S1), which was performed with 24-well microplates specific for microbe culture (IWAKI) as previously described^22^. The *E. coli* cells were cultured in eight wells, and eight tenfold serial dilutions, i.e., 10^1^~10^8^, were prepared with fresh medium. The microplates were incubated overnight in a microplate bioshaker (Deep Well Maximizer, Taitec) at 37°C, with rotation at 500 rpm. Serial transfer was performed at 12- or 24-h intervals, according to the growth rate. Only one of the eight wells (dilutions) showing growth in the early exponential phase (OD_600_ = 0.01-0.1) was selected and diluted into eight wells of a new microplate using eight dilution ratios. The cell culture selected daily for the following serial transfer was mixed with glycerol (15% v/v) and stored at −80°C for future analyses. Serial transfer was repeatedly performed for approximately 50 days. The evolutionary generation was calculated according to the following equation (Eq. 1).

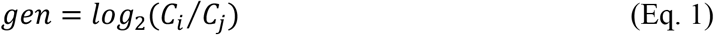

Here, *C_i_* and *C_j_* represent the OD_600_ of the cell culture that was used for serial transfer and the theoretical OD_600_ of the cell culture at the start of incubation. *C_j_* was calculated by dividing the OD_600_ that was used in the last transfer by the dilution rate. To benefit experimental replication, the cell cultures stored for the following assays were dispensed into 20 microtubes in small aliquots (100 μL per tube), which were used once, and the remainder was discarded, as previously described^55^.

### Growth fitness assay

The fitness was determined as the maximal growth rate, as previously reported^20^. In brief, the cell culture stocks were diluted 1,000-fold in fresh media (C0, C1~28) and were subsequently loaded into a 96-well microplate (Costar) in six wells at varied locations. The 96-well microplate was incubated in a plate reader (Epoch2, BioTek) with a rotation rate of 567 rpm at 37°C. The temporal growth of the *E. coli* cells was detected by measuring the absorbance at 600 nm, and readings were obtained at 30-min intervals for 48 h. The maximal growth rate was calculated according to the following equation (Eq. 2).

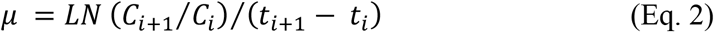

Here, *C_i_* and *C*_*i*+1_ represent the two reads of OD_600_ values at two `consecutive` time points of *t_i_* and *t*_*i*+1_ The growth fitness was the average of the five continuous growth rates that exhibited the largest mean and the smallest standard deviation during the temporal changes in growth rate, as previously reported^20^. A total of 2,220 growth curves were acquired, and the corresponding growth rates were calculated for the analysis (Table S5).

### Redox activity assay

A cell culture in the exponential phase of growth (OD_600_ = 0.01 ~ 0.3) was used for the assay. The cell culture was diluted with fresh medium at twelve dilution ratios from 1.75^0^ to 1.75^11^ in a final volume of 2 mL. Every 100 μL of the diluted cell culture was transferred to multiple wells in a 96-well microplate (Costar), in which 20 μL of CellTiter 96^®^ Aqueous One Solution Reagent (Promega) was added. The reduction of the tetrazolium compound in the reagents was measured with a microplate reader (Epoch2, BioTek) by determining the OD_490_ every 2 min for 30 min. The rate of reduction was calculated by linear regression of the temporal changes in OD_490_, i.e., the slope of the increase in OD_490_ over time (min). The redox activity was determined by dividing the rate of reduction by the OD_600_ of the cell culture. The mean of the multiple measurements (N=5) was used for the analysis.

### Niche space evaluation

The fitness dynamics (i.e., the fitness curve) across the concentration gradient of each chemical component were evaluated by curve fitting of a cubic polynomial with the following equation (Eq. 3).

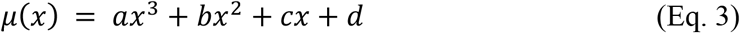

Here, *x* and *μ*(*x*) represent the concentration gradient of each chemical component and the growth rate under the corresponding conditions, respectively. *a*, *b, c* and *d* are the constants. The area under the fitted curve was calculated according to the following equation (Eq. 4).

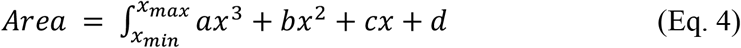

Here, *x_min_* and *x_max_* represent the minimum and maximum concentrations of each chemical component, respectively. The niche space (*S*) was evaluated by normalizing both the height and the width of the fitness curve with the following equation (Eq. 5).

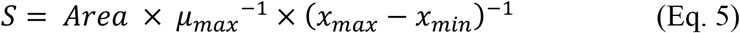

Here, *μ_max_* is the maximal growth rate across the concentration gradient. The niche broadness (*S_T_*) of the individual genome was determined as the sum of the niche spaces of the eight chemical components as follows (Eq. 6).

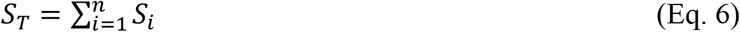

Here, *S_i_* and *n* indicate the niche space of each chemical component and the total number of chemical components, respectively.

### Genome resequencing and mutation analysis

The stored cell culture was inoculated into 4 mL of fresh M63 medium in a test tube and grown at 37°C with shaking at 200 rpm. Once cell growth reached the stationary phase (OD_600_ > 1.0), rifampicin was added to the culture at a final concentration of 300 μg/mL to stop genome replication initiation. After 3 h of culture with rifampicin, the cells were collected as previously reported^56^. Genomic DNA was extracted using a Wizard Genomic DNA Purification Kit (Promega) in accordance with the manufacturer’s instructions. The sequencing libraries were prepared using the Nextera XT DNA Sample Prep Kit (Illumina), and paired-end sequencing (300 bp × 2) was performed with the Illumina MiSeq platform. The sequencing reads were aligned to the *E. coli* W3110 reference genome (AP009048.1, GenBank), and the genome mutations were analysed with the Breseq pipeline (version 0.30.1)^57^. The fixed mutations (Table S3) were subsequently analysed for the temporal order of accumulation during evolution. The raw data set was deposited in the DDBJ Sequence Read Archive under accession number DRA011629.

### Sanger sequencing and single-colony isolation

The genomic region of approximately 300-600 kb centred on the position of the mutation was amplified by PCR with PrimeSTAR HS DNA Polymerase (TaKaRa Bio) and the corresponding primers (Table S4). Amplicons were purified using a MinElute PCR Purification Kit (Qiagen), and Sanger sequencing was conducted by Eurofins Genomics K. K. (Tokyo, Japan). The resulting electropherogram was analysed using Sequence Scanner Software v2.0 (Thermo Fisher Scientific), and the ratio of the mutants within the cell population was calculated according to the peak values, as described previously ^58^. Stored cell cultures with an interval of ~100 generations were analysed to identify the heterogeneity of the cell population. Single-colony isolation was performed from the heterogeneous population to isolate the homogeneous mutants. The cell culture was spread on LB agar plates, and 10~30 single colonies per plate were subjected to Sanger sequencing. The colonies of the homogeneous mutant were stored for the fitness assay as described above.

## Authors’ contributions

MK and IN performed the experiments, MK and BWY analysed the data and drafted the manuscript, BWY conceived the research and rewrote the paper, and all authors approved the final manuscript.

## Acknowledgements

We thank NBRP for providing the *E. coli* strains carrying the wild-type and reduced genomes (KHK collection). This work was supported by the JSPS KAKENHI Grant-in-Aid for Scientific Research (B) (grant number 19H03215 (to BWY)).

