## Supplementary figures and tables for "Local adaptation mediated niche expansion in correlation with genetic richness"

Supplementary figures (Figures S1~S7) pp.2~8
Supplementary tables (Table S1~S4) pp.9~12

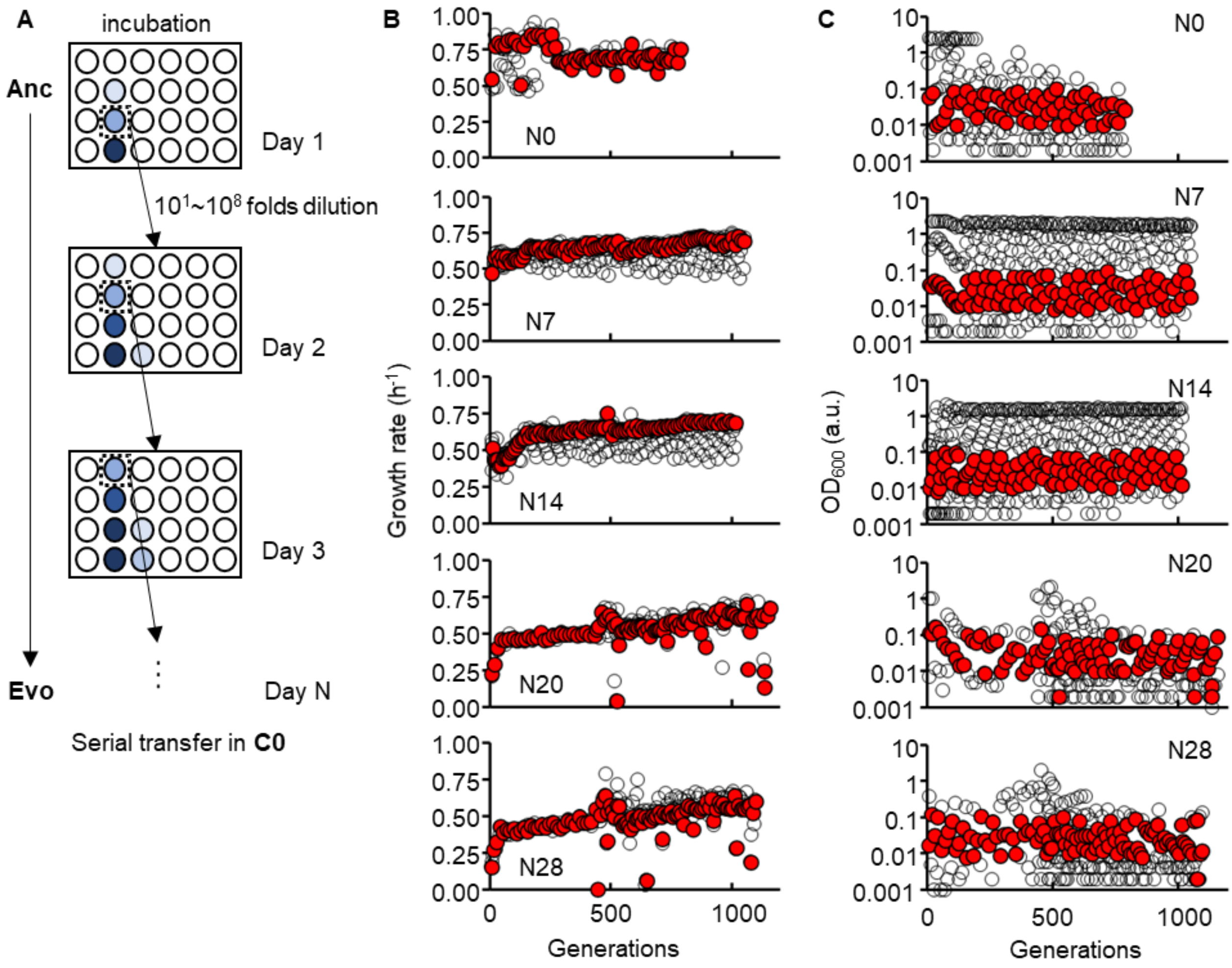


**Figure S1 Experimental evolution. A.** Schematic drawing of the evolution experiment. **B.** Temporal changes in growth rate during evolution. The genomes are indicated. The grey and red circles indicate the cultures with varied dilution rates and the one out of eight cultures selected for serial transfer, respectively. **C.** Temporal changes in $OD_{600}$ during the evolution. The grey and red circles indicate the eight cultures and the one selected for serial transfer, respectively. The $OD_{600}$ of the selected cultures roughly ranged from 0.01 to 0.1.

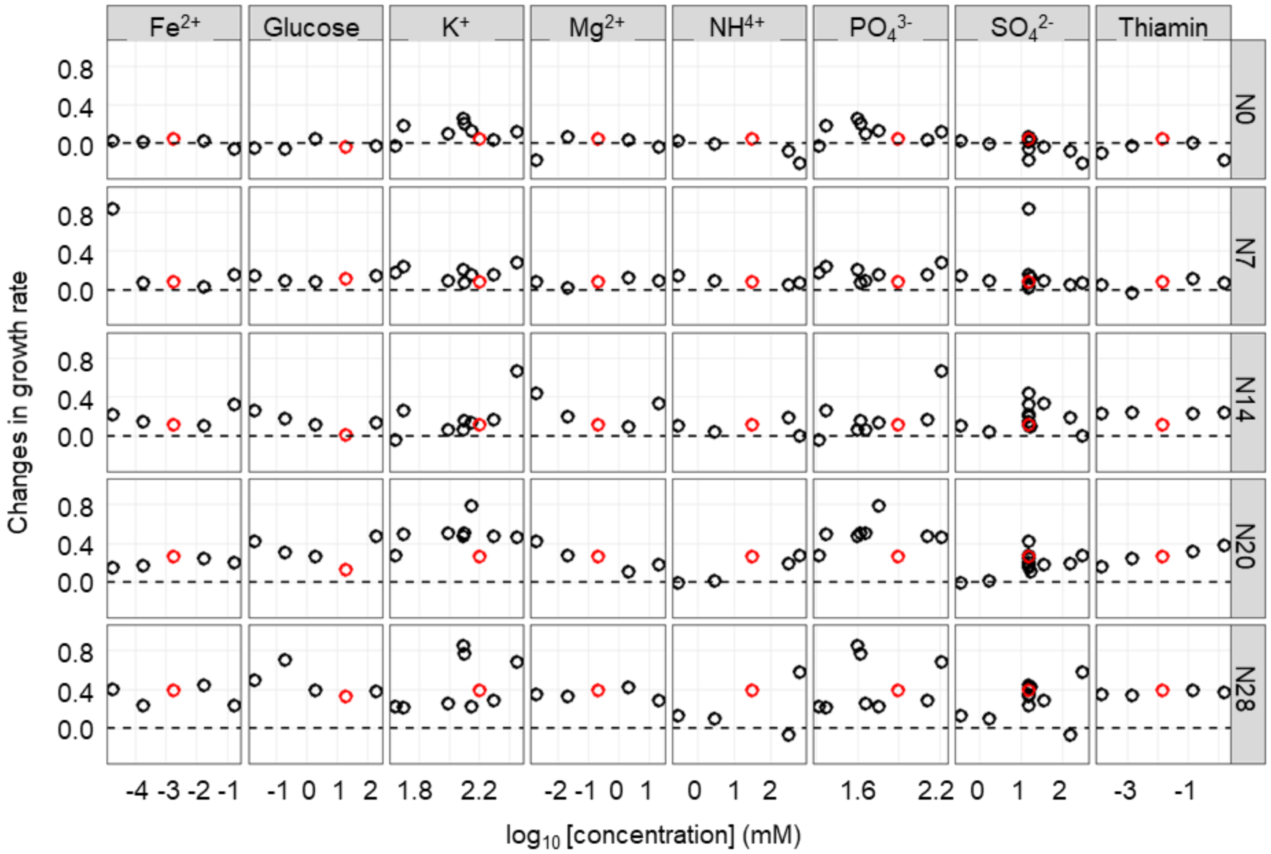


**Figure S2 Changes in growth rate across the concentration gradient of individual chemical components.** The changes in fitness, which were acquired by subtracting the mean growth rate of Anc from that of Evo, are shown. The changes in C0 and C1~C28 are indicated in red and black circles, respectively. The concentrations of the chemical components are shown on a logarithmic scale. The wild-type and reduced genomes are indicated as N0 and N7~N28, respectively.

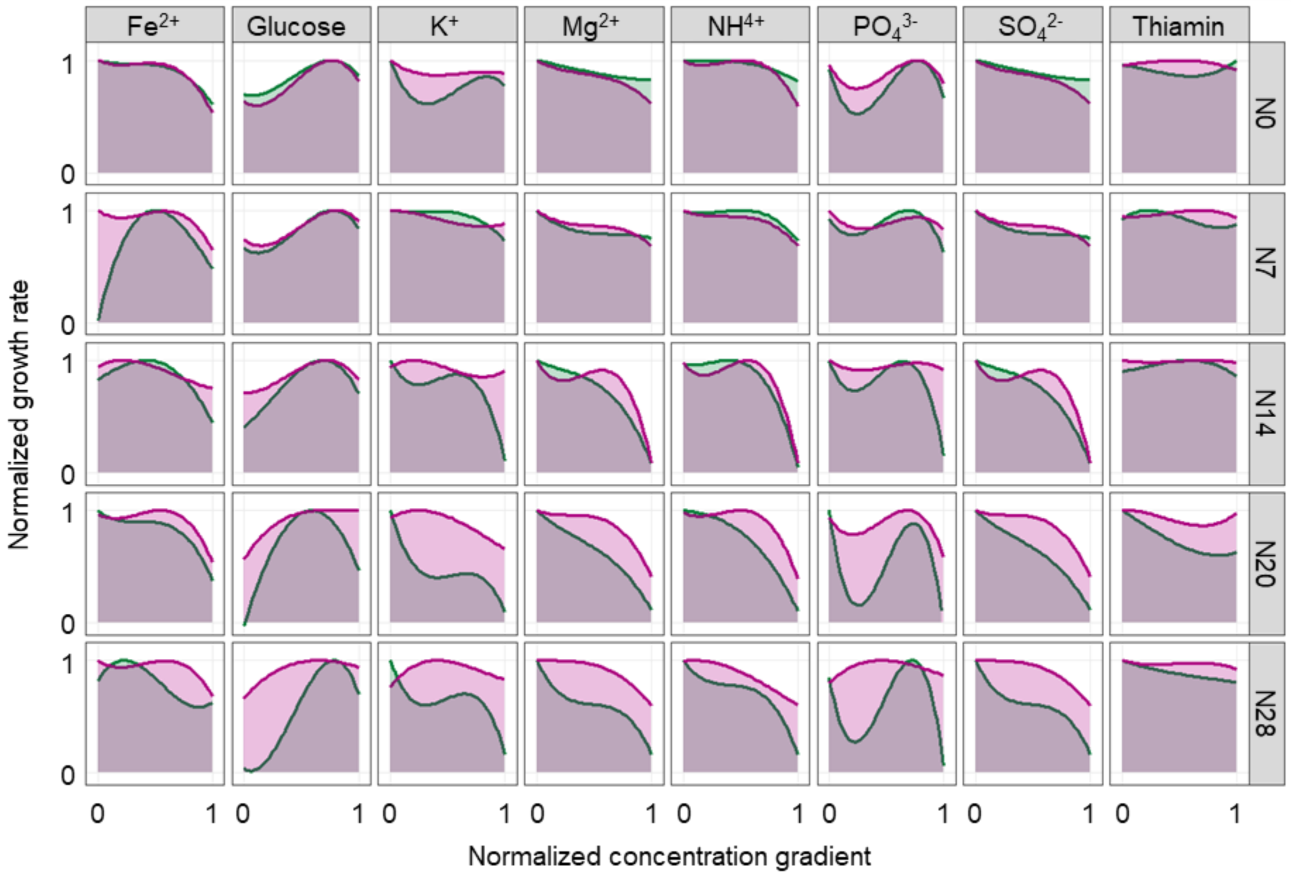


**Figure S3 Niche spaces of Ancs and Evos.** The fitness curves are normalized, in which both the concentration gradient and the growth rates are rescaled within one unit. The transparent green and purple areas indicate the niche spaces of Ancs and Evos, respectively.

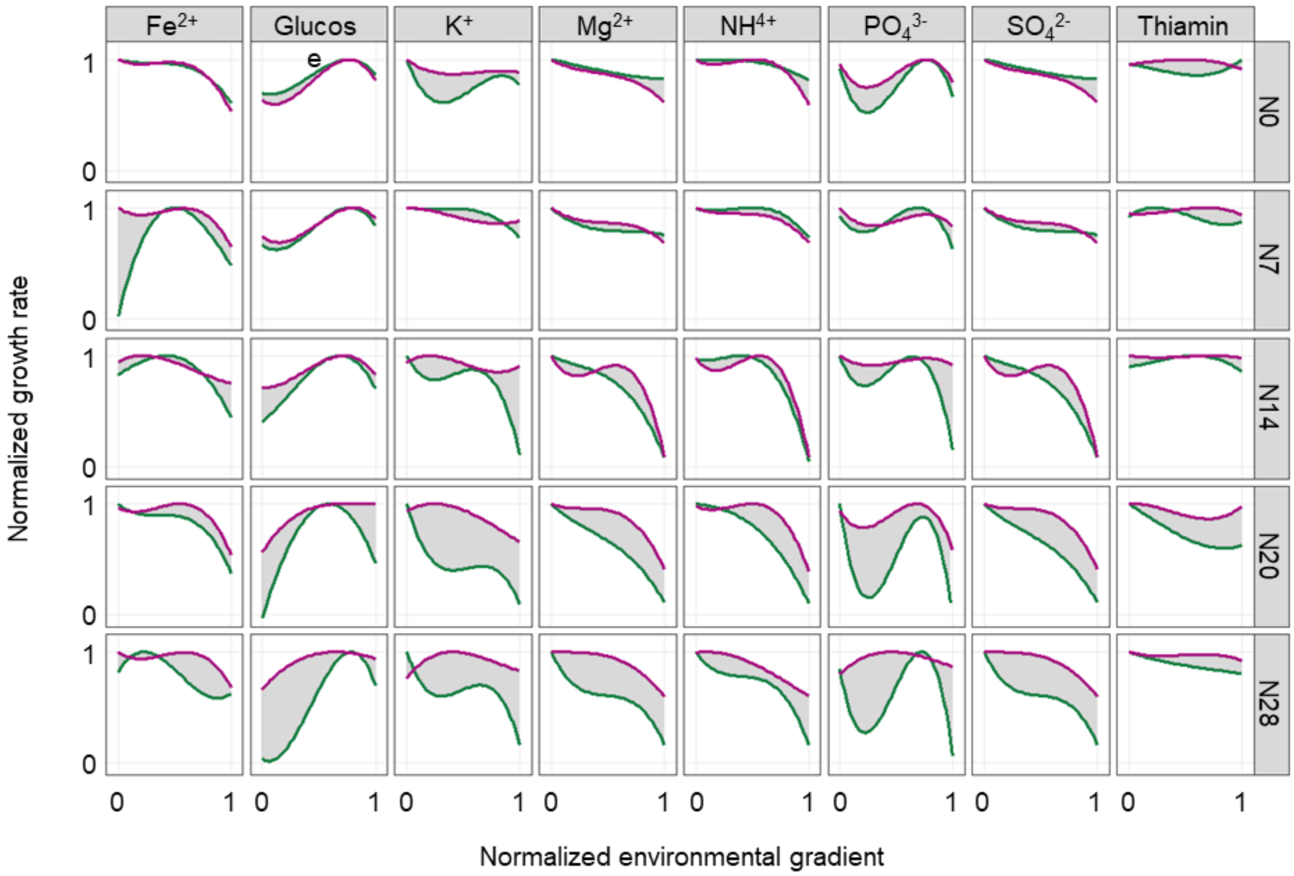


**Figure S4 Changes in niche spaces between Ancs and Evos.** The green and purple lines indicate the fitness curves of Ancs and Evos, respectively. The shadowed areas represent the changes in niche spaces between Ancs and Evos.

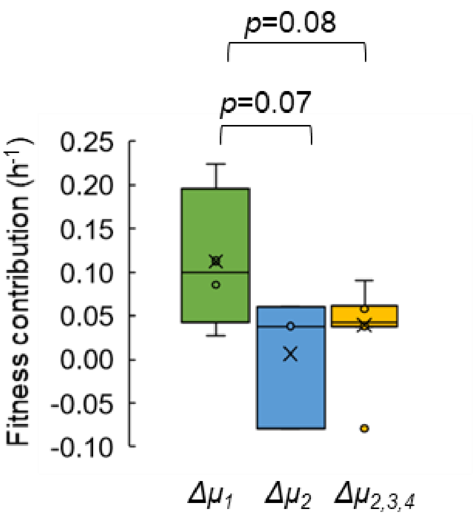


**Figure S5 Boxplot of the changes in growth rates caused by mutation accumulation.** Green, blue, and yellow represent the changes in growth rates caused by the first mutations, the second mutations, and the second, third and fourth mutations in the four genomes (N7, N14, N20 and N28), respectively. The statistical significance is indicated.

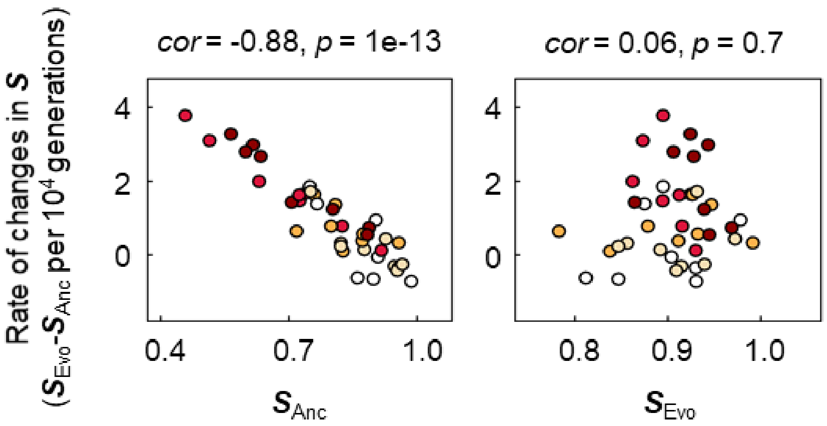


**Figure S6 Relationships of $S_{Anc}$ and $S_{Evo}$ to the rate of changes in niche spaces.** The rates of changes in the niche space per generation are plotted against $S_{Anc}$ and $S_{Evo}$ in the left and right panels, respectively. The colour variation from white to dark red represents the five different genomes of N0~N28. The Spearman rank correlation coefficients and statistical significance are indicated.

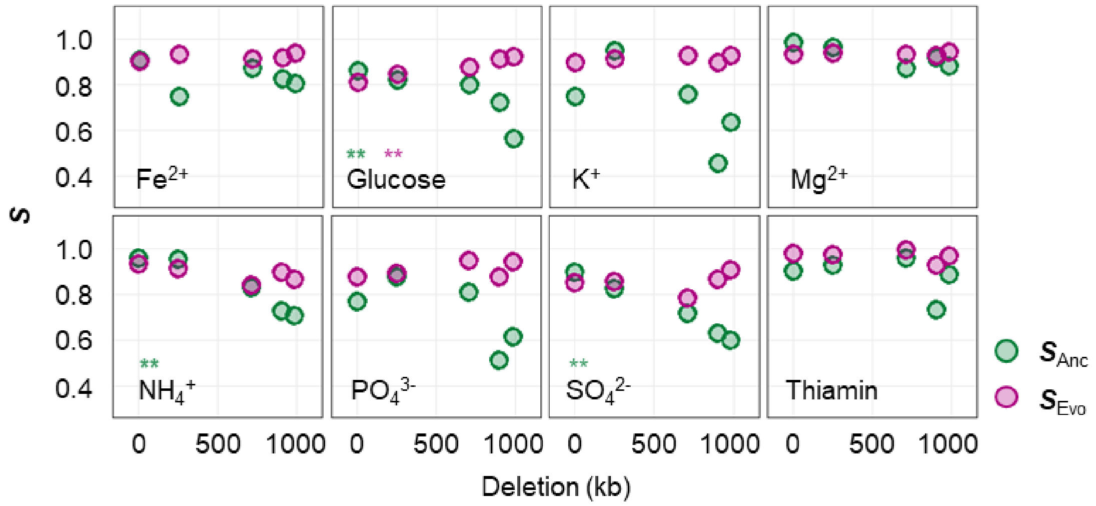


**Figure S7 Relationships between niche space and genome reduction.** The niche spaces of the five genomes are plotted against the lengths of genomic deletion with respect to the eight chemical components (niches). Green and purple indicate Ancs and Evos, respectively. The chemical components (niches) and the statistical significance of the Spearman rank correlation are indicated. Asterisks represent statistical significance ($^{*}$, $p < 0.05$; $^{**}$, $p < 0.01$).

| Genomes | Strains | Genome size (bp) | Deletion (bp) | Deleted ratio (%) |
|---|---|---|---|---|
| N0 | W3110S | 4,646,332 | 0 | 0 |
| N7 | 07stepAW | 4,395,545 | 250,787 | 5.4 |
| N14 | 14stepAγW | 3,936,788 | 709,544 | 15.3 |
| N20 | 20stepCW | 3,747,380 | 898,952 | 19.3 |
| N28 | MGF-01W | 3,663,949 | 982,383 | 21.1 |

**Table S1 Genomes used in the present study.** Genome IDs are the labels of the wild-type (N0) and the reduced (N7, N14, N20, N28) genomes used in the study. Strain names are the names of the *E. coli* strains carrying the corresponding genomes, which were deposited in the National BioResource Project (NBRP). Genome size, deletion and deletion ratio indicate the full length of the genome, the deleted length and its ratio compared to the wild-type genome, respectively.

| Environmental gradient | $KH_2PO_4$ (mM) | $K_2HPO_4$ (mM) | $FeSO_4$ (mM) | Thiamine (mM) | $(NH_4)_2SO_4$ (mM) | $MgSO_4$ (mM) | Glucose (mM) |
|---|---|---|---|---|---|---|---|
| C0 | 38.2 | 61.6 | 0.0018 | 0.0148 | 15.3 | 0.203 | 20 |
| C1 | 38.2 | 61.6 | 0.000018 | 0.0148 | 15.3 | 0.203 | 20 |
| C2 | 38.2 | 61.6 | 0.00018 | 0.0148 | 15.3 | 0.203 | 20 |
| C3 | 38.2 | 61.6 | 0.018 | 0.0148 | 15.3 | 0.203 | 20 |
| C4 | 38.2 | 61.6 | 0.18 | 0.0148 | 15.3 | 0.203 | 20 |
| C5 | 38.2 | 61.6 | 0.0018 | 0.000148 | 15.3 | 0.203 | 20 |
| C6 | 38.2 | 61.6 | 0.0018 | 0.00148 | 15.3 | 0.203 | 20 |
| C7 | 38.2 | 61.6 | 0.0018 | 0.148 | 15.3 | 0.203 | 20 |
| C8 | 38.2 | 61.6 | 0.0018 | 1.48 | 15.3 | 0.203 | 20 |
| C9 | 38.2 | 61.6 | 0.0018 | 0.0148 | 15.3 | 0.00203 | 20 |
| C10 | 38.2 | 61.6 | 0.0018 | 0.0148 | 15.3 | 0.0203 | 20 |
| C11 | 38.2 | 61.6 | 0.0018 | 0.0148 | 15.3 | 2.03 | 20 |
| C12 | 38.2 | 61.6 | 0.0018 | 0.0148 | 15.3 | 20.3 | 20 |
| C13 | 38.2 | 61.6 | 0.0018 | 0.0148 | 15.3 | 0.203 | 0.02 |
| C14 | 38.2 | 61.6 | 0.0018 | 0.0148 | 15.3 | 0.203 | 0.2 |
| C15 | 38.2 | 61.6 | 0.0018 | 0.0148 | 15.3 | 0.203 | 2 |
| C16 | 38.2 | 61.6 | 0.0018 | 0.0148 | 15.3 | 0.203 | 200 |
| C17 | 38.2 | 61.6 | 0.0018 | 0.0148 | 0.153 | 0.203 | 20 |
| C18 | 38.2 | 61.6 | 0.0018 | 0.0148 | 1.53 | 0.203 | 20 |
| C19 | 38.2 | 61.6 | 0.0018 | 0.0148 | 153 | 0.203 | 20 |
| C20 | 38.2 | 61.6 | 0.0018 | 0.0148 | 306 | 0.203 | 20 |
| C21 | 1.91 | 61.6 | 0.0018 | 0.0148 | 15.3 | 0.203 | 20 |
| C22 | 3.82 | 61.6 | 0.0018 | 0.0148 | 15.3 | 0.203 | 20 |
| C23 | 19.1 | 61.6 | 0.0018 | 0.0148 | 15.3 | 0.203 | 20 |
| C24 | 76.4 | 61.6 | 0.0018 | 0.0148 | 15.3 | 0.203 | 20 |
| C25 | 38.2 | 3.08 | 0.0018 | 0.0148 | 15.3 | 0.203 | 20 |
| C26 | 38.2 | 6.16 | 0.0018 | 0.0148 | 15.3 | 0.203 | 20 |
| C27 | 38.2 | 30.8 | 0.0018 | 0.0148 | 15.3 | 0.203 | 20 |
| C28 | 38.2 | 123.2 | 0.0018 | 0.0148 | 15.3 | 0.203 | 20 |

**Table S2 Compositions of the 29 medium combinations.** The chemical compounds used to generate the medium combinations (C) are indicated. The final concentrations of the seven compounds in the media are shown.

| Strain | Changes in DNA | Changes in AA | Gene name | Description |
|---|---|---|---|---|
| N7 | G→A | intergenic (+267/-28) | ycfH / ptsG | predicted metallodependent hydrolase / fused glucose-specific PTS enzyme IIBC components |
| | G→A | intergenic (-59/-156) | fliE / fliF | flagellar basal-body component / flagellar basal-body MS-ring and collar protein |
| | C→A | G139W (GGG→TGG) | trkH | potassium transporter |
| | Δ1 bp | coding (417/993 nt) | rbsR | DNA-binding transcriptional repressor |
| N14 | G→A | R220H (CGT→CAT) | secF | SecYEG protein translocase auxillary subunit |
| | A→C | L18F (TTA→TTC) | malX | fused maltose and glucose-specific PTS enzyme IIBC components |
| | C→T | G359D (GGT→GAT) | cyaA | adenylate cyclase |
| | Δ2 bp | coding (708/2022 nt) | rep | DNA helicase and single-stranded DNA-dependent ATPase |
| N20 | C→T | G235S (GGT→AGT) | sapD | predicted antimicrobial peptide transporter subunit |
| | T→G | H40Q (CAT→CAG) | ynbC | predicted hydrolase |
| | G→A | N14N (AAC→AAT) | yggR | predicted transporter |
| N28 | Δ1 bp | coding (828/876 nt) | cysW | sulfate/thiosulfate transporter subunit |
| | C→T | G255D (GGC→GAC) | kgtP | alpha-ketoglutarate transporter |
| | C→A | D570E (GAC→GAA) | rpoD | RNA polymerase, sigma 70 (sigma D) factor |

**Table S3 Gene mutations.** The genome mutations fixed in the evolved populations of the reduced genomes (N7, N14, N20 and N28) are summarized. No mutation was detected in the evolved N0. Nt and Aa represent nucleotides and amino acids, respectively. The slash indicates the noncoding region between the two genes indicated. The description indicates the function of the gene product.

| ID | Target | Sequence (5' to 3') |
|---|---|---|
| ycfH_ptsG_F | *ycfH / ptsG* | GTGCAACTTCTCCAATGATCTG |
| ycfH_ptsG_R | | GTTATTGGTAAAGCCGAGGGC |
| fliEF_F | *fliE / fliF* | CTAATGGTCGGTTGCGGCAG |
| fliEF_R | | GGAACCGGCAACAATCAATG |
| trkH_F | *trkH* | GGCATCCATTCCGGCAAACC |
| trkH_R | | GATAGTGGTGCTGTTCTGGAC |
| rbsR_F | *rbsR* | CACCAGTGACTTCATAGCCATC |
| rbsR_R | | CACTGCCAGTACCAATCCTTTC |
| secF_F | *secF* | CGCTGCTGTCTATCCTCGTG |
| secF_R | | GGAAACACCGATAAGCATGGTC |
| malX_F | *malX* | CCATGCAGATGACCTACTCC |
| malX_R | | CGATACAGAACATGACAGGCAG |
| cyaA_F | *cyaA* | CTTCAAACGCGGCATACAGC |
| cyaA_R | | GGAGCGTGTTACTGAATACCTG |
| rep_F | *rep* | CGTGCGGGTTATTGGCGATC |
| rep_R | | GTATGATGCACACCTGAAAGC |
| sapD_F | *sapD* | CCAGACGACAACCAATCGGTAAC |
| sapD_R | | CTGCTGATTGCTGACGAACCG |
| ynbC_F | *ynbC* | CAGGCATTATTAGCGGGTTTG |
| ynbC_R | | CAAGAGATGGGCTATAACCACG |
| yggR_F | *yggR* | GGCAGTGCGAAGGTAACAGC |
| yggR_R | | GTTACGCCCAGTTCCTGAAAG |
| cysW_F | *cysW* | GTGCCGTGGAAGCGAATATG |
| cysW_R | | GTCGCTGCCGTTACAGATTG |
| kgtP_F | *kgtP* | GCCAGCGAACCGAAACATAAC |
| kgtP_R | | GTTGTGGCGTTGTGGTTACG |
| rpoD_F | *rpoD* | GGAAGACAAGATCCGCAAAGTG |
| rpoD_R | | CTGGAGAAACTGGTTGAAGCG |

**Table S4 Primers for Sanger sequencing.** The sequences of the primers used to examine the population heterogeneity of the mutations determined by genome resequencing.
